# Predicting biomass of resident kōkopu (*Galaxias*) populations using local habitat composition

**DOI:** 10.1101/2021.12.16.472917

**Authors:** Ben R. J. Crichton, Michael J.H. Hickford, Angus R. McIntosh, David R. Schiel

## Abstract

With the global decline of freshwater fishes, quantifying the body size-specific habitat use of vulnerable species is crucial for accurately evaluating population health, identifying the effects of anthropogenic stressors, and directing effective habitat restoration. Populations of New Zealand’s endemic kōkopu species (*Galaxias fasciatus, G. argenteus*, and *G. postvectis*) have declined substantially over the last century in response to anthropogenic stressors, including habitat loss and fragmentation, invasive species, and over-exploitation. Despite well-understood habitat associations, key within-habitat features driving the reach-scale biomass of small and large kōkopu remain unclear. Here, we investigated whether the total biomass of small (≤ 90 mm) and large (> 90 mm) kōkopu was associated with total pool area, average pool depth, total bank cover, average substrate size, and average forest canopy cover across fifty-seven 50 m reaches. These features were selected because generally pool habitats are productive feeding areas, bank cover and substrate interstices are important refuges, and forest cover provides greater food availability. Because kōkopu are nocturnal, populations were sampled with removal at night using headlamps and hand-nets until reaches were visually depleted. Using Akaike’s information criterion, it was found that increases in large kōkopu biomass were most parsimoniously explained by greater pool area and bank cover, whereas increases in small kōkopu biomass were best explained by low bank cover and greater average forest cover. This study demonstrated the importance of considering the ontogenetic shift in species’ habitat use and provided an effective modelling approach for quantifying the size-specific habitat use of these stream-dwelling fish.

## Introduction

Given the widespread decline of freshwater fishes [1], it is crucial to quantify which habitats are used during all stages of a species’ life cycle so that population health can be accurately evaluated, effects of anthropogenic stressors can be tested, and successful rehabilitation measures implemented [2, 3]. Anthropogenic stressors such as pollution, habitat fragmentation and degradation, introduced species, river regulation, and over-exploitation have contributed to a substantial decline in riverine fish populations over the last century [4, 5]. Unfortunately, many statistical models used for studying the effects of anthropogenic stressors on populations are inaccurate due to being calibrated using only a fraction of the habitats used by a species [6]. Without accurate models relating body size and specific habitats, population assessments may be biased, which could lead to ineffective management decisions and unsuccessful, wasteful, or even harmful restoration efforts by excluding important microhabitats such as spawning sites or nursery grounds [7, 8].

Influential habitat variables that often determine the habitat selection of stream-dwelling fish include water velocity, in-stream refuges, and overhanging vegetation [9-11]. Pools are often preferentially used microhabitats for freshwater fish because they have slower water velocities, which typically reduce an individual’s energetic expenditure [12, 13] while improving feeding efficiency [14, 15]. In-stream cover, such as undercut banks, root-wads, debris dams, and interstices between large substratum are important refuges that many fish rely on to minimise the risk of predation and the impacts of physical disturbances [16, 17]. Additionally, overhanging vegetation, such as riparian vegetation or forest canopy cover, is linked to a stream’s primary productivity and plays a crucial role in providing terrestrial subsidies, in-stream cover, and hydrological stability [18, 19]. Therefore, these habitat features are likely influential determinants of habitat selection during at least one stage of the lifecycle of stream-dwelling fishes.

The importance of specific habitat features on habitat selection is often strongly determined by body size [20]. In freshwater fishes, variation in size-related habitat selection is typically due to individual selection of microhabitats that maximise energy gain and minimise energy expenditure or increase survival [21-23]. Because microhabitat selection is strongly linked to individual fitness, species may rely on several distinct microhabitats to support different size-classes [23, 24]. For species where different size-classes inhabit the same local environment, it is vital that restoration efforts incorporate potential ontogenetic shifts in size-specific microhabitat requirements to account for all size-classes in an ecosystem. This is especially important for species that exhibit intraspecific or intra-family competitive hierarchies, because inferior individuals may avoid preferred habitats when dominant congeners are present [25]. Social competitive hierarchies in freshwater fish often follow a size-related structure; large dominant individuals monopolise key feeding habitats and smaller individuals are displaced to less advantageous habitats [26, 27]. Therefore, understanding how abiotic and biotic influences affect the habitat use of distinct size-classes is essential to obtain a robust evaluation of population habitat use.

New Zealand’s endemic banded kōkopu (*Galaxias fasciatus*), giant kōkopu (*G. argenteus*), and shortjaw kōkopu (*G. postvectis*), hereafter collectively referred to as ‘kōkopu’, are diadromous fishes that inhabit the same stream environments during all but their larval life stage. Over the last century, kōkopu have undergone considerable declines in response to a combination of habitat loss, migratory barriers, introduced species, and fishing pressure [28-30]. The loss and degradation of adult habitats through activities including drainage schemes, land-use change, and deforestation are thought to be the biggest drivers of decline in kōkopu [29, 31]. Migratory barriers inhibit upstream dispersal to compatible habitats [32, 33] and introduced species like trout alter kōkopu habitat selection through predation and competitive exclusion [34, 35]. Post-larval kōkopu are also harvested in the culturally, recreationally, and commercially important whitebait fishery [36]. Despite population declines, it is unknown how these anthropogenic stressors specifically alter kōkopu populations due to the lack of accurate size-specific habitat models.

Although size-specific habitat models have not been developed for kōkopu, there is a thorough understanding of general habitat preferences [37]. Greater kōkopu densities are often associated with the availability of slow-flowing pools because kōkopu are opportunistic, mostly nocturnal predators, that rely on the transport of aquatic and terrestrial invertebrates into pools from fast-flowing upstream habitats [29, 38]. Banded and shortjaw kōkopu are forest specialists, rarely inhabiting streams without forest canopies, but giant kōkopu also inhabit estuaries, swamps, or ponds [37, 39]. Each species depends on refuge areas for secure diurnal resting, predator escapement, and shelter from flood events [40]. Despite having slightly different habitat preferences, the kōkopu species commonly co-occur and share similar environmental requirements (i.e., diet and habitat use), which likely indicates that each species should be influenced similarly by changes to habitat composition from anthropogenic stressors within stream environments.

Even though juvenile and adult kōkopu occupy the same local environments, individual microhabitat selection is strongly determined by the presence of larger conspecifics or congenerics [41]. For example, small giant kōkopu minimise agonistic interactions with larger dominant conspecifics that control large pools at night by feeding during the day or by occupying alternative microhabitats at night [42]. Similarly, large banded kōkopu prefer deep, slow-flowing pools, with coarse substratum, whereas smaller individuals are likely displaced into shallow pools with faster water velocities and finer substratum [43]. Although size-related kōkopu microhabitat segregation [27, 44, 45] and the influence of habitat composition on total kōkopu biomass [11, 25] are understood, how within-habitat characteristics influence the reach-scale biomass of small and large kōkopu separately is unknown. Such information would provide a more comprehensive and accurate description of kōkopu habitat requirements that could be used to improve habitat restoration efforts. Additionally, by understanding how small and large kōkopu are influenced by local environments, while all other influential environmental variables are being controlled for, a standardised prediction of likely kōkopu biomass based solely on local habitat characteristics can be obtained. These standardised estimates will allow the isolation and accurate testing of how individual environmental manipulations including dispersal barriers, introduced predators, fishing pressure, or habitat restoration efforts affect kōkopu populations by removing habitat-related biases.

To examine how kōkopu size-classes respond to habitat composition, all three kōkopu species were studied as one overall ‘population’ because body size is likely the key driver of habitat use, they commonly co-occur, have similar habitat requirements, and abide by intra-family competitive group behaviours [27, 46]. These factors likely mean that one species’ position in a stream could be used by either of the other species if it was vacant. A size-class break-point of 90 mm (total length) was used to examine how small and large kōkopu respond to habitat composition. This break-point was selected because banded kōkopu and giant kōkopu are approximately one year old at this size and begin to compete for territory when >90 mm [47-49]. Equivalent studies have not been completed with shortjaw kōkopu, but they were pooled into the same size-class groups for consistency. The similar ecological and physiological characteristics between kōkopu species strongly suggest that the compiled grouping of species will allow an accurate investigation of size-related habitat selection without species-specific biases.

We aimed to identify habitat features that influence the biomass of small and large kōkopu. Specifically, we used Akaike’s Information Criterion (AIC), an information theoretic approach [50], to evaluate a candidate set of *a priori* models to explain variation in small and large kōkopu biomass using local habitat features. To achieve this objective, kōkopu populations were surveyed across physically diverse stream reaches. We predicted that: (1) large kōkopu biomass would increase with pool area and depth, whereas small kōkopu biomass would decrease in such habitats, putatively due to larger fish competitively excluding smaller individuals within these key feeding areas; (2) large and small kōkopu biomass would increase with increasing bank cover and substrate size due to both providing refuges to all size-classes; and (3) both large and small kōkopu biomass would increase with forest canopy cover due to it likely providing greater food availability.

## Methods

### Study Sites

To investigate which habitat features are most strongly associated with reach-scale kōkopu biomass, three 50 m reaches were sampled within each of 19 streams on the West Coast of New Zealand’s South Island during May and June 2021. Local topographic maps, site visits, and databases, such as Freshwater Ecosystems of New Zealand (FENZ; [51]) and the New Zealand Freshwater Fish Database (NZFFD; [52]), were used to select streams that contained kōkopu and that had no fish passage barriers or trout presence. All streams were open to whitebait fishing because unfished streams were limited. Physically diverse streams that included a wide range of habitat compositions were selected to provide a robust understanding of how individual habitat variables influenced kōkopu biomass. Sampling took place within two months to minimise seasonal differences in kōkopu biomass.

### Habitat survey

Study reaches began and ended at riffles, which acted as minor fish barriers between reaches, were located in areas with minimal surface turbulence or natural visual-obstruction deposits (i.e., foam or fine debris collections), and were no deeper than 1.5 m. Habitat surveys, completed during daylight hours, involved measuring the area and average depth of pools, availability of in-stream bank cover, average substrate size, and percentage cover of forest canopy within each reach. Forest cover was measured at approximately eight locations within each reach using a spherical crown densiometer [53] while standing in the middle of the waterway and facing upstream. In-stream bank cover was recorded by measuring the perimeter of root wads, undercut banks, or debris dams accessible to fish. Pool area was calculated using:

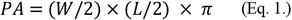

where *PA* is pool area (m^2^), *W* is the maximum width of the pool (m), and *L* is the maximum length of the pool (m). Average pool depth was calculated from ten depth measurements along the *W* axis. The average substrate size within each reach was calculated from approximately 60 stones randomly selected using a Wolman’s walk [54].

### Kōkopu biomass survey

The three 50 m reaches within each stream were sampled starting >1 h after sunset (using spotlighting) when kōkopu are active. Sampling consisted of counting kōkopu using a high-powered spotlight to scan the reach for fish in slow-flowing habitats [55]. This method has been used effectively for sampling kōkopu within wadeable streams at night [34, 56]. Alternative fish sampling methods such as electrofishing and trapping are generally less effective for surveying kōkopu because they sink when stunned, occupy deep bank cover during the day, and may not encounter traps due to having high pool fidelity at night [57-59]. The 1 h delay after sunset ensured that resident kōkopu had left their daytime refuges and moved into nocturnal foraging areas where they could be seen and captured. When spotted, kōkopu generally remained stationary and were caught using hand nets. All captured fish were placed into buckets of aerated stream water. When kōkopu were seen but not caught, the estimated length and species of the individual were recorded and noted as a ‘miss’. Reaches were sampled using successive depletion passes until fish were no longer observed. This required up to five passes and took around 1.5 h per reach. Captured kōkopu were anaesthetised in 20 mg/L of AQUI-S water-dispersible liquid anaesthetic to facilitate handling. The total length of each fish was measured on a wet measuring board (±1 mm) before being weighed (±0.01 g). After measurements were taken, fish were placed in buckets of fresh stream water to recover, and then returned to their area of capture. All procedures were approved by the University of Canterbury Animal Ethics Committee (permit number 2020/06R).

### Data analysis

Prior to analyses, large and small kōkopu biomass responses were square root transformed to meet assumptions of normality, and outliers (two large kōkopu responses and one small kōkopu response) were identified and removed using interquartile range criterion. Biomass measurements were used as a response instead of counts because kōkopu body mass varies substantially between individuals and is associated with available resources, whereas the association between fish counts and resource availability is also determined by competitive interactions [25]. Variance inflation factors (VIF) were calculated to ensure that there was no collinearity between predictors (i.e., VIF ≤ 4; [60]). Because all VIF values were low (VIF < 2.0) we proceeded with model selection.

To assess how local habitat composition influenced kōkopu biomass, a set of ecologically realistic *a priori* linear mixed-effects models, which included all combinations of the five habitat variables, was used to explain the biomass of each kōkopu size-class. Ecologically realistic interactions between habitat features were initially included, but later removed due to poor data spread creating unreliable results. Linear mixed-effects models were constructed using the ‘lme’ function (Package ‘nlme’; [61]) in R version 4.1.1 [62] and included a random factor for stream so that each of the three reaches nested within a stream could be independently used to examine how habitat composition influenced kōkopu biomass. By focusing on the reach-scale, more accurate and informative localised habitat-biomass relationships could be obtained.

An information theoretic approach, using Akaike’s information criterion corrected for small sample size (AIC_c_), was used to determine which candidate models explained variation in large and small kōkopu biomass most parsimoniously [50]. Each model’s AIC_c_ was subtracted from the lowest AIC_c_ to determine its ΔAIC_c_ [50]. Parsimonious models had ΔAIC_c_ values < 2 [63]. Conditional coefficient of determination (R^2^ _c_; proportion of variance explained by fixed and random effects) values were calculated for each parsimonious model to evaluate goodness-of-fit because AIC_c_ only ranks models relative to each other [64, 65]. The Akaike weight and R^2^ _c_ of parsimonious models were compared to select the most suitable model for explaining large and small kōkopu biomass.

Partial dependence plots, which show the independent effect of a single variable on the response by accounting for the average effects of all other variables in a model [66], were used to visually examine the independent effect of each habitat feature on the total biomass of large and small kōkopu. Using the ‘effects’ package [67], partial dependence plots were developed by extracting the independent effects of each variable within a linear mixed-effects model that included all five habitat features and a random factor for stream on the biomass of each size-class.

## Results

Large kōkopu biomass was explained parsimoniously (i.e., a ΔAICc < 2) by two models (Table 1). Predictors in the first model (1L) were total bank cover and pool area, while the second model (2L) also included total bank cover and pool area, but added average substrate size. Large kōkopu biomass was positively correlated with total pool area (R^2^_c_=0.19, F_1,53_=12.45, P<0.001; Fig. 1a), total bank cover (R^2^ _c_=0.26, F =18.75, P<0.001; Fig. 1c), and average substrate size (R^2^ _c_=0.11, F_1,53_ =6.79, P=0.012; Fig. 1d). However, there was no correlation between large kōkopu biomass and average pool depth (R^2^ _c_ <0.01, F_1,53_ =0.02, P=0.883; Fig. 1b) or forest cover (R^2^ _c_ <0.1, F_1,53_ = 0.45, P=0.505; Fig. 1e). Despite having fewer explanatory variables, model 1L explained just 4% less variation than model 2L (*R*^*2*^ _*c*_*=* 0.51 and 0.55, respectively; Table 1). Model 1L also better accounted for variation in large kōkopu biomass, as indicated by the Akaike weights of 0.21 and 0.16, respectively. This suggested that model 1L was the most suitable for predicting large kōkopu biomass. Table 2 details the summary statistics for model 1L.

**Table 1.** Top linear mixed-effects models (ΔAIC_c_ < 2) that explain variation in the total biomass of large and small kōkopu based on Akaike’s information criterion (AIC).

| Response | Model | Fixed effects | $AIC_c$ | $\Delta AIC_c$ | $w$ | $R^2_c$ |
| --- | --- | --- | --- | --- | --- | --- |
| Large kōkopu |  |  |  |  |  |  |
| $\sqrt{\text{Large kōkopu biomass}}$ | **1L** | **BC + PA** | **346.7** | **0.00** | **0.20** | **0.51** |
|  | 2L | BC + PA + SS | 347.1 | 0.46 | 0.16 | 0.55 |
| Small kōkopu |  |  |  |  |  |  |
| $\sqrt{\text{Small kōkopu biomass}}$ | **1S** | **FC + BC** | **203.4** | **0.00** | **0.20** | **0.58** |
|  | 2S | FC + BC + SS | 204.6 | 1.20 | 0.11 | 0.58 |
|  | 3S | FC + BC + PA | 205.1 | 1.78 | 0.08 | 0.61 |
|  | 4S | FC | 205.3 | 1.96 | 0.07 | 0.5 |

$AIC_c$ represents AIC values corrected for small sample size, delta $AIC_c$ ( $\Delta AIC_c$ ) is the difference in $AIC_c$ score between the highest ranked model and the candidate model, the Akaike weight ( $w$ ) is the probability that a particular model is the most parsimonious model among the candidate models, and $R^2_c$ is the conditional coefficient of determination. 'BC' is total bank cover (m), 'PA' is total pool area (m<sup>2</sup>), 'SS' is average substrate size (cm), and 'FC' is mean forest cover (%). Models are listed from lowest to highest $AIC_c$ score within each size-class. Bolded models were selected as the most suitable for predicting biomass within each size-class.

**Fig 1.**
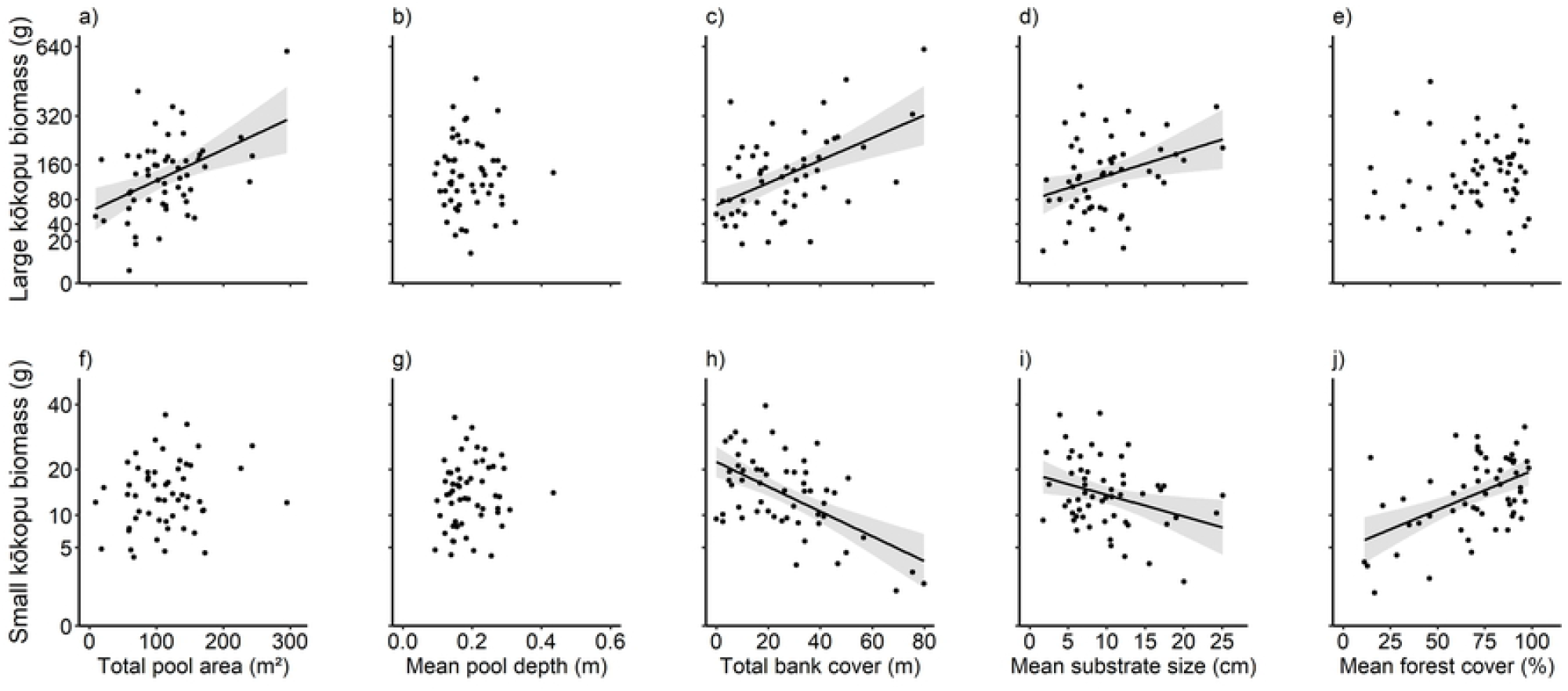
Relationships between habitat features and kōkopu biomass. Partial regression plots showing the independent effect of total pool area (a, f), mean pool depth (b, g), total bank cover (c, h), mean substrate size (d, i), and mean forest cover (e, j) on large (row 1), and small kōkopu biomass (row 2). Note that the Y-axis is not linear. Dots represent the total biomass of giant, banded, and shortjaw kōkopu in the size-class within each 50 m reach. Lines of best fit are shown where a significant correlation was found (P < 0.05) and error bands show 95% confidence intervals determined from model fits.

**Table 2.** Summary results of the fixed effects included in the linear mixed-effects models that most parsimoniously predict the total biomass of large and small kōkopu as identified in Table 1.

| Model | Fixed effects | Coefficient estimate (± SE) | P value |
| --- | --- | --- | --- |
| Large kōkopu |  |  |  |
| $\sqrt{\text{Large kōkopu biomass}} \sim \text{BC} + \text{PA}$ | Intercept | 5.016 (± 1.986) | 0.016 |
| | BC | 0.108 (± 0.049) | 0.036 |
| | PA | 0.031 (± 0.015) | 0.04 |
| Small kōkopu |  |  |  |
| $\sqrt{\text{Small kōkopu biomass}} \sim \text{FC} + \text{BC}$ | Intercept | 3.033 (± 0.756) | <0.001 |
| | FC | 0.020 (± 0.009) | 0.041 |
| | BC | -0.026 (± 0.012) | 0.035 |

‘BC’ is total bank cover (m), ‘PA’ is total pool area (m<sup>2</sup>), and ‘FC’ is mean forest cover (%).

In contrast to large kōkopu biomass, small kōkopu biomass was negatively correlated with bank cover (R^2^ _c_ =0.33, F_1,54_=27.05, P<0.001; Fig. 1h) and substrate size (R^2^ _c_ =0.11, F_1,54_=6.33, P=0.012; Fig. 1d), but positively correlated with forest cover (R^2^ _c_ =0.25, F_1,54_=18.64, P<0.001; Fig. 1j). Additionally, small kōkopu biomass was not correlated with pool area (R^2^ _c_ =0.03, F_1,54_=1.78, P=0.188; Fig. 1f) or pool depth (R^2^ _c_ =0.01, F_1,54_=0.78, P=0.382; Fig. 1g). Four models explained small kōkopu biomass parsimoniously (Table 1). The first model (1S), which included forest cover and bank cover, explained 3% less variation than the most explanatory model (3S), which also included pool area (*R*^*2*^ _*c*_ = 0.58 and 0.61, respectively). However, model 1S was 9% more likely to explain variation in small kōkopu biomass most parsimoniously than the second model (2S), which also included average substrate size (Akaike weights = 0.20 and 0.11, respectively). This suggested that model 1S was the most suitable for predicting small kōkopu biomass. Table 2 details the summary statistics for model 1S.

## Discussion

The quantification of body size with respect to specific habitat use is crucial for accurately identifying key habitats that support all life stages of a species and directing beneficial management and restoration efforts [68]. We aimed to identify key habitat features that influence the biomass of small and large kōkopu, and to create statistical models that can predict kōkopu biomass based on local habitat features while controlling for other influences. Our results indicate that small and large kōkopu have distinct habitat requirements, and the influence of habitat composition on biomass was not consistent between size-classes. By characterising the effects of local habitat composition on the biomass of small and large kōkopu separately, we provide a more comprehensive and accurate description of kōkopu habitat requirements.

Total pool area was a key habitat feature for explaining variation in large kōkopu biomass, whereas average pool depth had little influence. This indicates that large kōkopu can use most pool habitats, regardless of depth. Pool habitats are commonly associated with greater biomasses of large stream-dwelling fish like salmonids [69, 70]. Positive correlations between pool area and large fish biomass is expected, because although faster water velocities transport more drifting invertebrates downstream, slower flowing habitats such as pools promote greater feeding success by increasing strike efficiency and prey capture [23]. However, species like kōkopu and trout will maximise their net energy gain by occupying slow-flowing pools below fast-flowing reaches [2, 15, 71]. Unlike trout, which are predominantly diurnal visual predators, nocturnal galaxiids rely mainly on mechanical lateral line and olfactory sensory systems that work most effectively in slow-velocities [72, 73]. Therefore, similarly to other large stream fish, slow-flowing pools likely support greater large kōkopu biomass because they are profitable foraging areas.

Despite pool area being a key habitat requirement for large kōkopu, neither total pool area nor average pool depth influenced small kōkopu biomass. The lack of relationship between small kōkopu and pool habitat was likely caused by the greater biomass of large congeners, which were not restricted by pool depth, competitively displacing smaller individuals [45, 74]. Small kōkopu may also avoid pools because large individuals cannibalise smaller congeners [44, 75]. Similar relationships have been observed in drift-feeding cutthroat trout (*Oncorhynchus clarkii*); large fish occupy deep pools and smaller young-of-the-year inhabit shallower water [76]. However, in the absence of large conspecifics, small trout choose, and grow faster in, large pools over shallower water [26]. Although small kōkopu are likely displaced into less profitable foraging areas, they still select habitats with the lowest available velocity [43]. This suggests that slow-flowing pools may be included in the fundamental niche of small kōkopu, but biotic interactions with larger predators result in these areas falling outside of their realised niche.

In addition to pool habitat, in-stream refuges and large substratum were important habitat features that were positively associated with large kōkopu biomass. However, unlike their larger congeners, these features were negatively associated with small kōkopu biomass. Despite hypothesising that small kōkopu would also use these features for refuges, our results show that large kōkopu dominate these areas, suggesting they competitively displace smaller individuals from them. Similarly to kōkopu, in-stream cover is thought to be the most important habitat feature influencing juvenile and adult salmonid habitat selection [77]. However, most experimental studies that have added wood to streams have found that juvenile and adult salmonids respond positively [78]. It is important to consider that habitat structure can also increase predation risk by providing predator habitat [79]. In addition to large kōkopu, longfin eels (*Anguilla dieffenbachii*) also benefit from greater bank cover availability, which could also lead to small kōkopu avoiding these microhabitats [17]. Although there are contrasting accounts of preferred in-stream cover, adult banded kōkopu and giant kōkopu will readily use alternative bank cover when preferred cover is scarce [37]. This suggests that compiling various forms of bank cover into one variable is acceptable for kōkopu habitat-biomass modelling. In-stream cover is likely the most influential habitat feature on kōkopu biomass due to the strong conflicting effects on small and large kōkopu biomass.

Unlike pool area and in-stream cover, forest cover was not associated with large kōkopu biomass. This was unexpected because forested streams generally provide important terrestrially-derived food subsidies that can support greater fish biomass and contribute up to half the annual energy budget of some drift-feeding species [80, 81]. However, we sampled in autumn when terrestrial subsidies substantially reduce seasonally [81]. Terrestrial invertebrates are an essential food resource for banded kōkopu, making up 75% of their diet by number, and 89% by weight [58]. Importantly, our method of surveying forest canopy cover within reaches using a densitometer may not accurately represent the availability of terrestrial food resources because it measures the canopy immediately over the reach, whereas resources can be sourced from further upstream or from lower-growing riparian vegetation. Overall, forest cover is not locally important in explaining large kōkopu biomass.

In contrast to large kōkopu, forest cover was the only habitat feature that was positively associated with small kōkopu biomass. This was somewhat expected because banded kōkopu post-larvae migrate in greater abundances into streams that drain catchments with greater indigenous forest cover [36]. McDowall [82] hypothesised that kōkopu post-larvae may use warmer water temperatures to identify more-forested catchments in contrast to cooler streams that are derived from glaciers and mountainous regions. It is unclear to what extent small kōkopu benefit from terrestrially-derived food subsidies because their small gape size may inhibit the capture of larger terrestrial invertebrates [83]. Additionally, because small kōkopu are displaced competitively from key feeding areas such as pools, smaller fish would have less access to terrestrial invertebrates. However, forested streams can support much greater densities of mayflies, stoneflies, and caddisflies that are more suitable prey for fish with a smaller gape size [84]. The size-specific importance of stream shading could be attributed to smaller kōkopu being more at risk of predation from visually feeding avian predators such as kingfishers (*Todiramphus sanctus*) due to being displaced from daytime refuges. In less shaded shallow streams, small cutthroat trout were more susceptible to visual avian predators than large trout because of predator gape-limitations [85]. Furthermore, shade was particularly important when in-stream cover was limited [85]. This indicates that small kōkopu may occupy reaches with greater forest canopy cover to reduce the likelihood of predation rather than for terrestrial inputs.

The absence of mutually benefitting habitat features on small and large kōkopu biomass indicates that there is no key feature that can be used to benefit all of the life stages of kōkopu, and that habitat restoration efforts will need to consider small and large kōkopu habitats concurrently. Because of these conflicts, it is important to identify which habitat compositions provide the greatest benefits to the population of reproductively valuable adults over time [86]. If juvenile habitats are limited or degraded, adult populations may become limited by recruits [87]. However, if an adult population typically has excess recruits and is limited by habitat then the most beneficial management decisions could be to prioritise adult habitats. Often, a balance of adult and juvenile habitat requirements must be incorporated into management restoration to benefit the overall population. For example, gravel augmentation is a key tool used for restoring salmonid spawning and incubation grounds [88]. However, a conflict arose when adult Chinook salmon (*Oncorhynchus tshawytscha*) preferentially spawned in fine gravels where embryo survival was least likely [88]. Therefore, it was suggested that intermediate sized gravels would maximize overall reproductive success across both spawning and incubation life stages. Comparatively, despite being associated with a decrease in small kōkopu biomass, there would likely be greater conservation benefits in adding in-stream refuges into reaches with habitat-limited adult kōkopu populations and excess juveniles, due to adults being reproductively valuable. Further research is required to investigate which balance of juvenile and adult habitats provides the greatest benefits to kōkopu populations.

In conclusion, this study demonstrates the importance of examining size-related habitat use when identifying key habitats that support species, and provides a detailed and effective modelling approach for predicting small and large size-classes of stream fish using simple habitat measurements. We showed that large kōkopu biomass was best explained by a combination of total pool area and bank cover availability, whereas small kōkopu biomass was best explained by a combination of total bank cover and average forest cover. With this enhanced understanding of how kōkopu size-classes are influenced by their local environments, we can obtain a standardised prediction of likely kōkopu biomass based on local habitat characteristics. These standardised predictions can be used to isolate and accurately test how anthropogenic stressors affect populations of these declining endemic kōkopu species [86]. Modelling techniques such as those presented in this study will likely be a crucial tool used in conserving freshwater fish species by effectively evaluating population distributions and densities, streamlining habitat restoration efforts, and mitigating anthropogenic stressors [89].

## Acknowledgements

We thank the Marine Ecology Research Group for field assistance, and the New Zealand Ministry of Business, Innovation and Employment, Department of Conservation and University of Canterbury for support. All sampling was approved by the University of Canterbury Animal Ethics Committee (2020/06R).

